# Fluorescence Lifetime: Beating the IRF and interpulse window

**DOI:** 10.1101/2022.09.08.507224

**Authors:** Mohamadreza Fazel, Alexander Vallmitjana, Lorenzo Scipioni, Enrico Gratton, Michelle A. Digman, Steve Pressé

## Abstract

Fluorescence lifetime imaging (FLIM) has been essential in capturing spatial distributions of chemical species across cellular environments employing pulsed illumination confocal setups. However, quantitative interpretation of lifetime data continues to face critical challenges. For instance, fluorescent species with known *in vitro* excited state lifetimes may split into multiple species with unique lifetimes when introduced into complex living environments. What is more, mixtures of species, that may be both endogenous and introduced into the sample, may exhibit; 1) very similar lifetimes; as well as 2) wide ranges of lifetimes including lifetimes shorter than the instrumental response function (IRF) or whose duration may be long enough to be comparable to the interpulse window. By contrast, existing methods of analysis are optimized for well-separated and intermediate lifetimes. Here we broaden the applicability of fluorescence lifetime analysis by simultaneously treating unknown mixtures of arbitrary lifetimes– outside the intermediate, goldilocks, zone–for data drawn from a single confocal spot leveraging the tools of Bayesian nonparametrics (BNP). We benchmark our algorithm, termed BNP-lifetime analysis of BNP-LA, using a range of synthetic and experimental data. Moreover, we show that the BNP-LA method can distinguish and deduce lifetimes using photon counts as small as 500.

## Introduction

Amidst a number of fluorescence microscopy techniques^1–8^, fluorescence lifetime imaging microscopy (FLIM) has extensively contributed to our understanding of sub-cellular structures and processes^9–16^. In FLIM experiments within a biological sample, multiple biomolecules may be labeled with unique fluorophores characterized by different lifetimes ^17–20^. To deduce how these labels are spatially distributed, a single (confocal) spot within the sample is exposed to either modulated^21,22^ or pulsed^23,24^ excitation with the excitation spot eventually scanned across the sample. Here, we focus on pulsed illumination since it provides time-stamped photon arrivals (and helps reduce phototoxicity). A train of such illumination pulses is shown in Fig. 1a.

**Figure 1:**
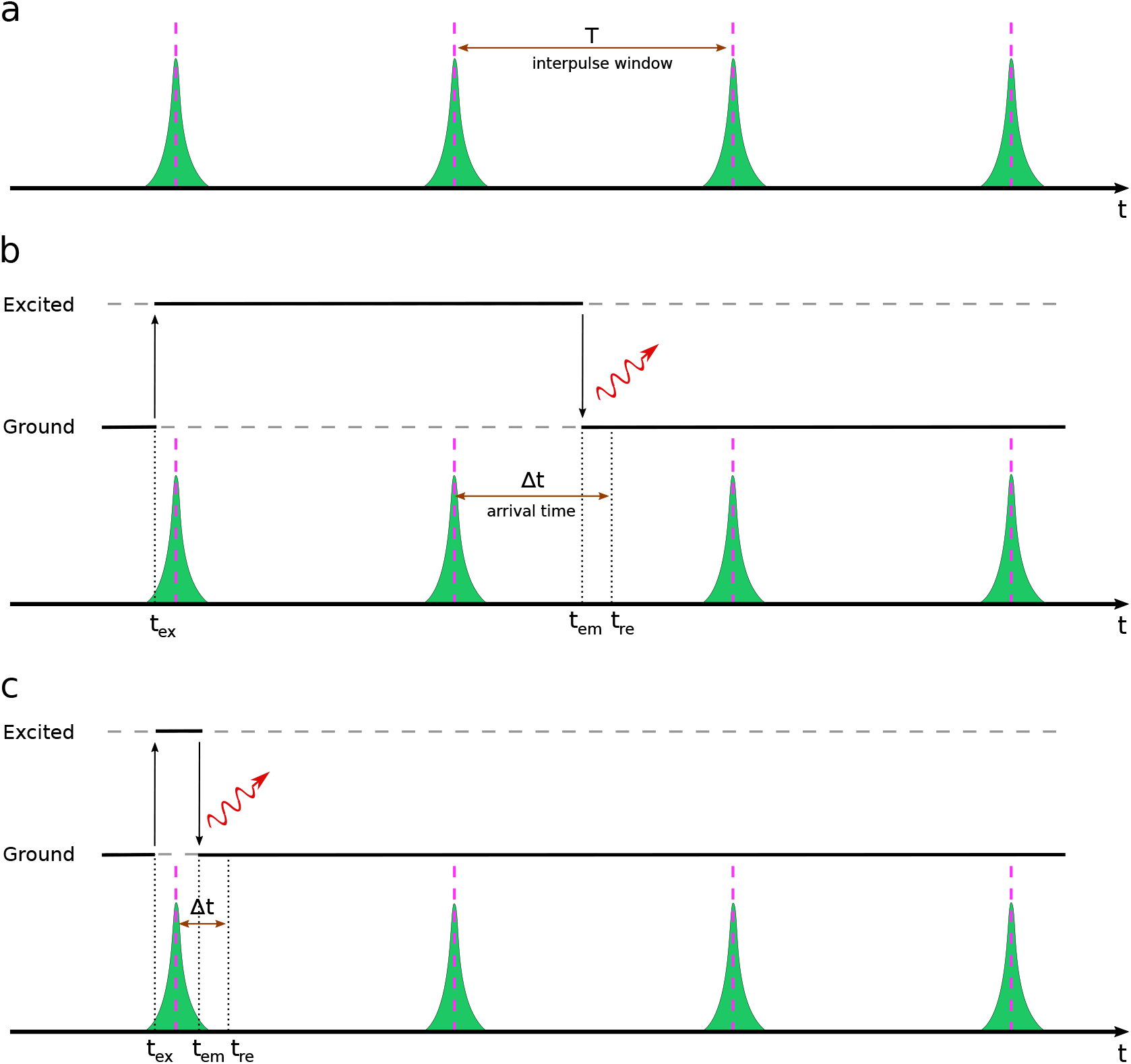
A cartoon representation of laser pulses, designated by green spikes, and fluorophore excitation and emission. (a) A train of laser pulses with interpulse window *T*. The pink dashed lines represent the pulse centers. (b) Fluorophores can be excited during laser pulses and may emit photons after multiple pulses due to long excited state lifetimes as compared to the interpulse windows. Indeed, even lifetimes on par with the interpulse time can appear after the subsequent pulse with probability *e*^*−*1^. (c) For fluorophores with lifetimes shorter than the IRF, an excited fluorophore might emit photons even before the excitation pulse is complete. Here, *t*_ex_, *t*_em_ and *t*_re_, respectively, stand for excitation, emission times and the recorded photon arrival time. The difference between emission and recorded times arises from the stochastic delay in detectors which, combined with the finite breadth of the laser pulse, is termed the IRF.

Once excited by a pulse, fluorophores emit photons whose arrival times at the detector are recorded and used to infer lifetimes and corresponding photon ratios across species for each spot^25–27^. By photon ratios we mean the probability (weight) that any given photon be emitted by each species within the confocal area probed. The photon ratio is itself related to the product of concentration of the species and excitation cross-section.

To deduce weights (photon ratios) and lifetimes present from photon arrival time data, analysis methods employ either model free techniques, such as phasors^26,28^ and deep learning^29,30^, or model based techniques, such as least-squares^31,32^, compressed sensing^33^, maximum likelihood^34,35^, and Bayesian methods^27,36–39^.

However, existing analysis methods are optimized for two well-separated lifetimes typically longer than the instrument response function (IRF) (see Fig. 1) but otherwise much shorter than the interpulse window. This regime of lifetimes can be difficult to control *in vivo* as lifetimes invariably drift in response to the local environment chemistry whose composition may further split apparent single lifetimes into multiple different lifetimes^40–42^. In addition, existing lifetime analysis methods, starting from the single spot/pixel, face several other key challenges including: 1) requiring the number of lifetime components as input otherwise often truncated for simplicity to two species^29,30,33–38^; 2) require high photon budgets due to information averaging arising from data pre-processing, *e*.*g*., data binning^31,32,43^; and 3) provide full uncertainty over the estimated parameters originating from unavoidable sources of stochasticity including random excitation times introduced by the IRF’s finite breadth and exponential waiting times for excited state lifetimes^29–35^.

To address these challenges, we begin by considering photon arrival times. These photon arrival times are essentially a mixture of temporal data points generated from multiple different sources, namely, fluorophore species, characterized by their lifetimes. As such, mathematically, the output of a pulsed excitation experiment may be conceptualized as generating data drawn from a mixture model where the ultimate goal of an analysis method would be to classify the arrival times into multiple categories corresponding to the underlying fluorophore species. More broadly, such classification tasks fall within the purview of clustering algorithms. For instance, K-means^44^ is perhaps the simplest and most popular clustering algorithm classifying a set of input data points into a given number, *K*, of clusters.

However, as the number of lifetime components is inherently unknown in photon arrival analysis, we need to evoke more sophisticated clustering algorithms.

To be precise, to correctly propagate inherent uncertainties, we work within a Bayesian paradigm where our inference is informed by sources of uncertainty including intrinsic stochasticity in the photon arrival times, finite breadth of IRF, and finite interpulse time. Moreover, we further specialize into working within a Bayesian nonparametric (BNP) paradigm to accomodate the unknown number of fluorophore species.

In particular, within BNPs, we leverage Dirichlet processes^45–49^ to allow inference over the number of species warranted by the data while rigorously propagating uncertainty from all the existing sources throughout the problem.

The Dirichlet process formally allows us to place priors on an infinite number of putative species that could be warranted by the data^45–49^. As we will see, as we collect data, weights associated to species contributing to the data will increase while the weights ascribed to other species will reduce to negligible values; see Fig. 2.

**Figure 2:**
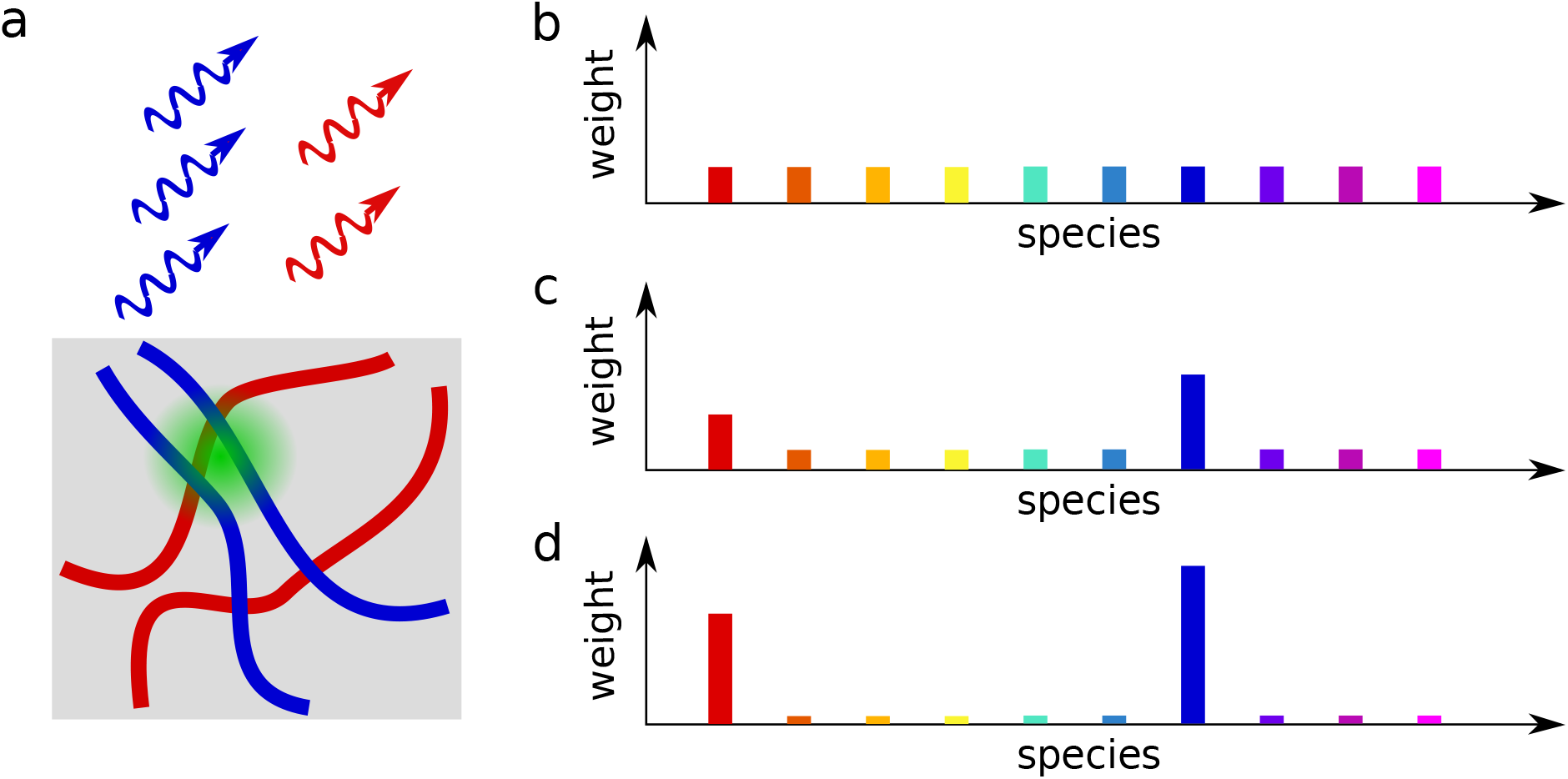
The Dirichlet process for lifetime analysis. (a) A spot within a sample is illuminated with a green laser which, in turn, leads to photons from red and blue fluorophore species staining different structures within the sample. The set of collected photon arrival times from this experiment is modeled with a Dirichlet process with the first ten species, represented by 10 different colors, shown. (b) When no photon arrival times have yet been collected, the weights ascribed to all species coincide with the prior value (often it is reasonable to assume uniform); (c) When 500 photon arrival times have been collected, the weights start differing from the nominal prior value and, in this cartoon, the blue and red species gain more weights; (d) At 5K photon arrival times most of weights are ascribed to the blue and red species while the rest tend toward negligible values.

Here, building upon our previous work^27,39^, we propose a BNP lifetime analysis (BNP-LA) method. This method leverages the Dirichlet process along with accurate likelihood model, informed by features such as IRF and pulsed excitation (see Fig. 1b-c), to simultaneously addresses all the following challenges: it is capable of dealing with a broad range of lifetimes ranging from values smaller than the IRF width to comparable to the interpulse time while addressing the challenges 1)-3) above.

Before moving onto the results, a word on nomenclature is warranted. “Species” here is defined as it normally is in the FLIM literature; *i*.*e*., as exponential components^1,2^. We thus inherit all advantages and disadvantages of this definition. This nomenclature is historically motivated by the fact that many fluorophore lifetime histograms are beautifully fit to a single exponential^36,39^. As such, we may be (incorrectly) inclined to assume that a species is a chemically distinct molecule. Indeed, we need to be careful as there exist cases where the lifetime may differ from exponential^50–52^ and be, say, bi-exponential. This is the case where two radiative pathways are available for de-excitation. In this case, the literature would define these as two “species”.

## Results

Our BNP-LA method’s main objective is to learn the lifetimes and their corresponding weights given a set of photon arrival times. As the BNP-LA method operates within the Bayesian framework, to learn these parameters we work with a posterior, which is proportional to the product of the likelihood and priors over these parameters (see Methods Section). However, our nonparametric posterior does not attain a standard form and we cannot deal with that analytically. Therefore, we develop a numerical strategy to efficiently sample our posterior (see Methods Section). The results presented in this section are thus histograms of samples drawn from the BNP-LA posterior.

Here, we use both synthetic and experimental data to evaluate the performance of our BNP-LA analysis package. We first use synthetic data to benchmark our method against: 1) a decreasing interpulse window (see Fig. 3) where photon detections occurring after pulses following the one inducing excitation become increasingly probable; 2) multiple lifetimes smaller than the width of IRF distribution, and lifetimes with sub-nanosecond differences (see Figs. 4-5); 3) photon counts (see Fig. 5); 4) a range of different weights associated to species due to variations in photon counts across species (see Fig. 6); and 5) more than two lifetimes (see Fig. 7).

**Figure 3:**
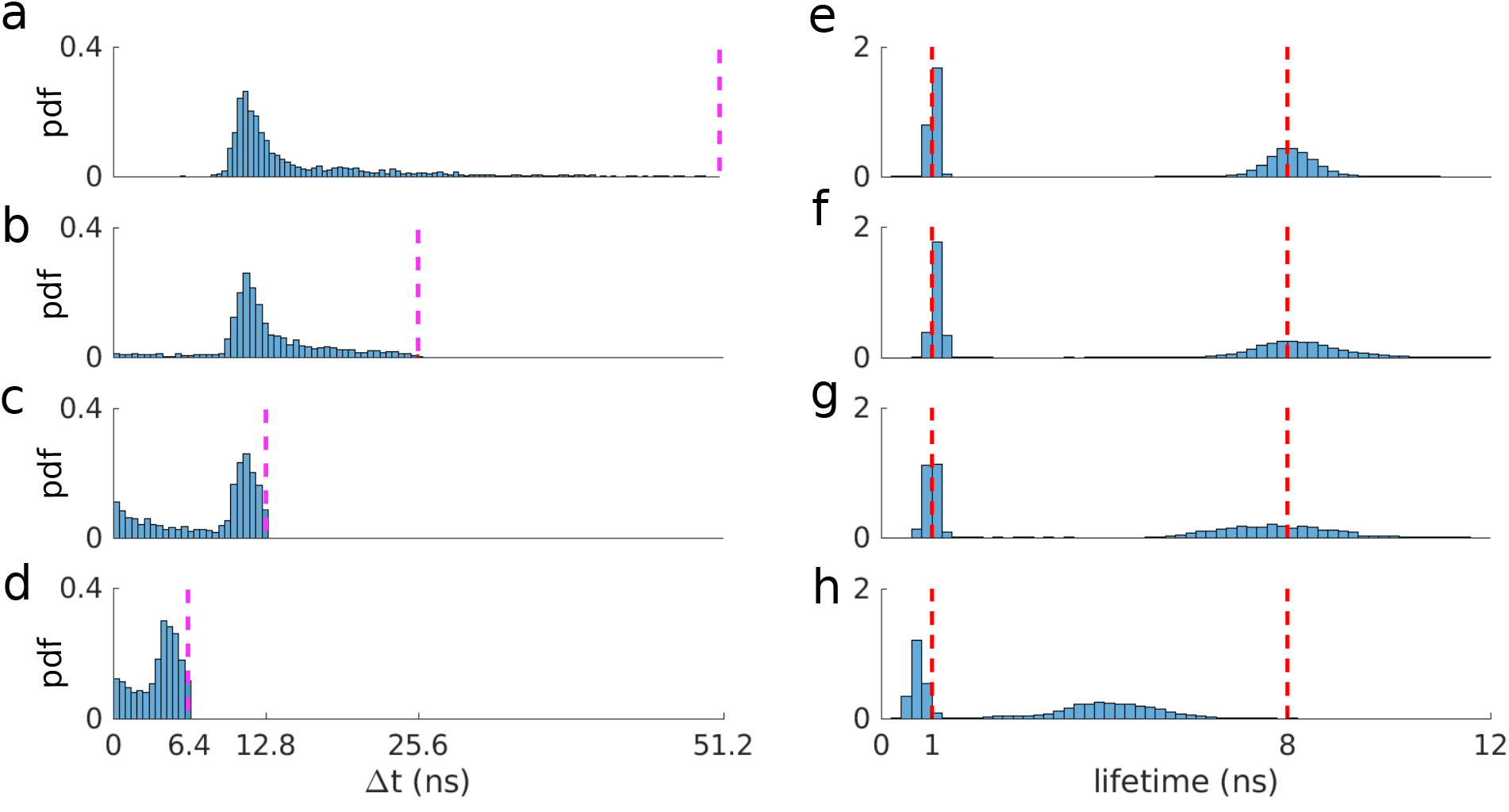
Interpulse window effect on lifetime estimation. (a-d) Histograms of photon arrival data generated with two lifetimes of 1 ns and 8 ns and interpulse times of 51.2, 25.6, 12.8 and 6.4 ns, respectively. Pink dashed lines represent the interpulse window. PDF stands for probability density function which is obtained by normalizing the area under histograms to unity. (e-h) Marginal posterior of lifetimes corresponding to each generated data. Red dashed lines represent ground truths. We retain the same convention throughout the manuscript.

**Figure 4:**
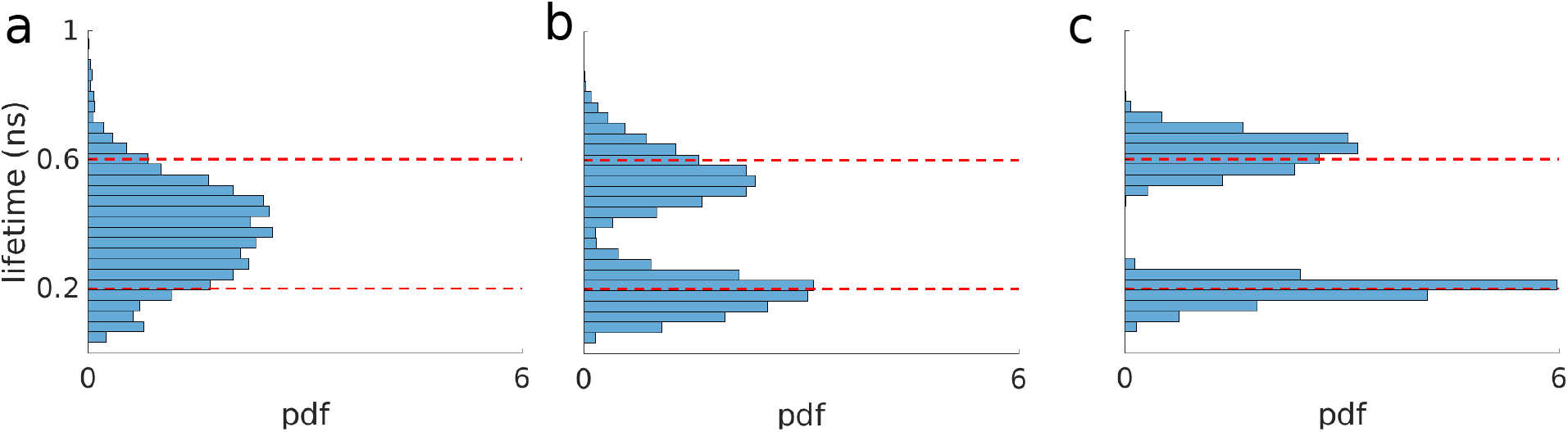
Two lifetimes smaller than the IRF width(*σ*_IRF_ = 0.66 ns). (a-c) Marginal posterior of lifetime with 500, 1K and 5K photons, respectively.

**Figure 5:**
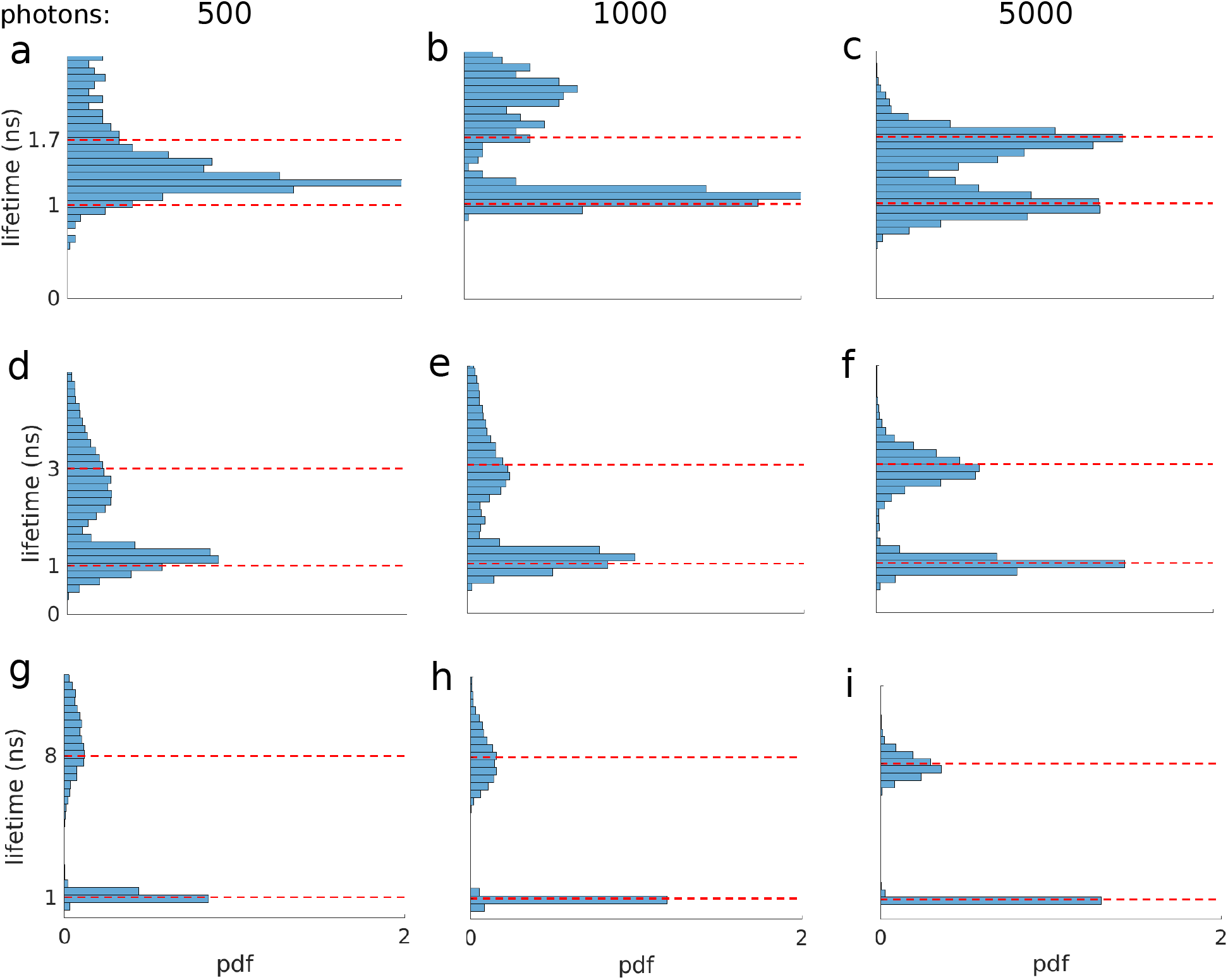
Robustness test against lifetimes and photon counts. (a-c) Marginal posteriors of lifetimes for 1 ns and 1.7 ns with sub-nanosecond difference using different photon counts. (d-f) Marginal posterior of lifetimes of 1 ns and 3 ns using different photon counts. (g-i) Marginal posterior of lifetimes of 1 ns and 8 ns (close to interpulse window) using different photon counts.

**Figure 6:**
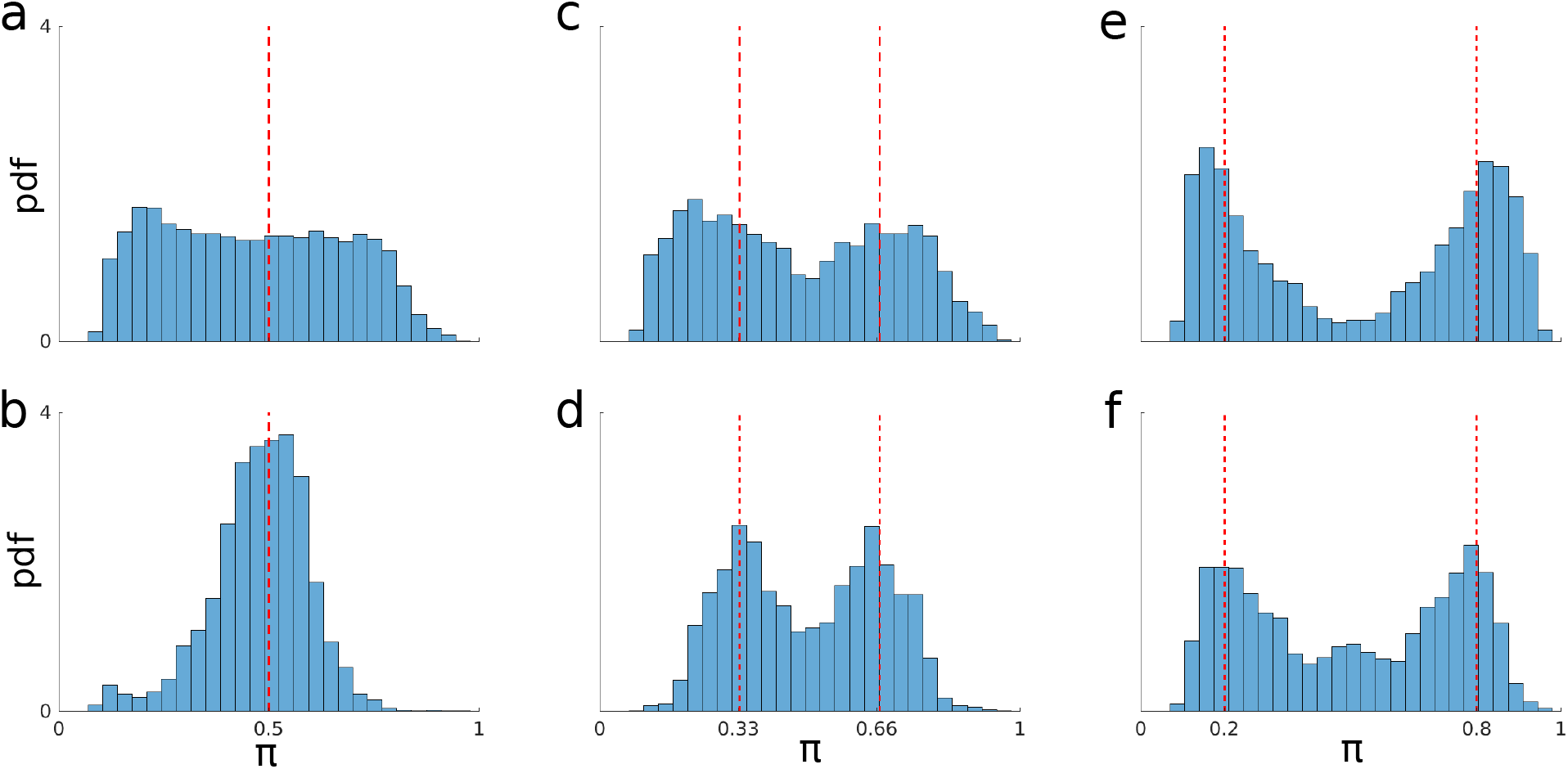
Robustness test against lifetime weights, shown by *π*, and photon counts. (a-b) Marginal posterior for the two weights found (when the ground truth is for 1/2 each) when 1K and 5K total photon counts are considered in the analysis, respectively. (c-d) Similar to above except for when the ground truth of the weights is 1/3 and 2/3. (e-f) Same as above excet for weights of 1/5 and 4/5.

**Figure 7:**
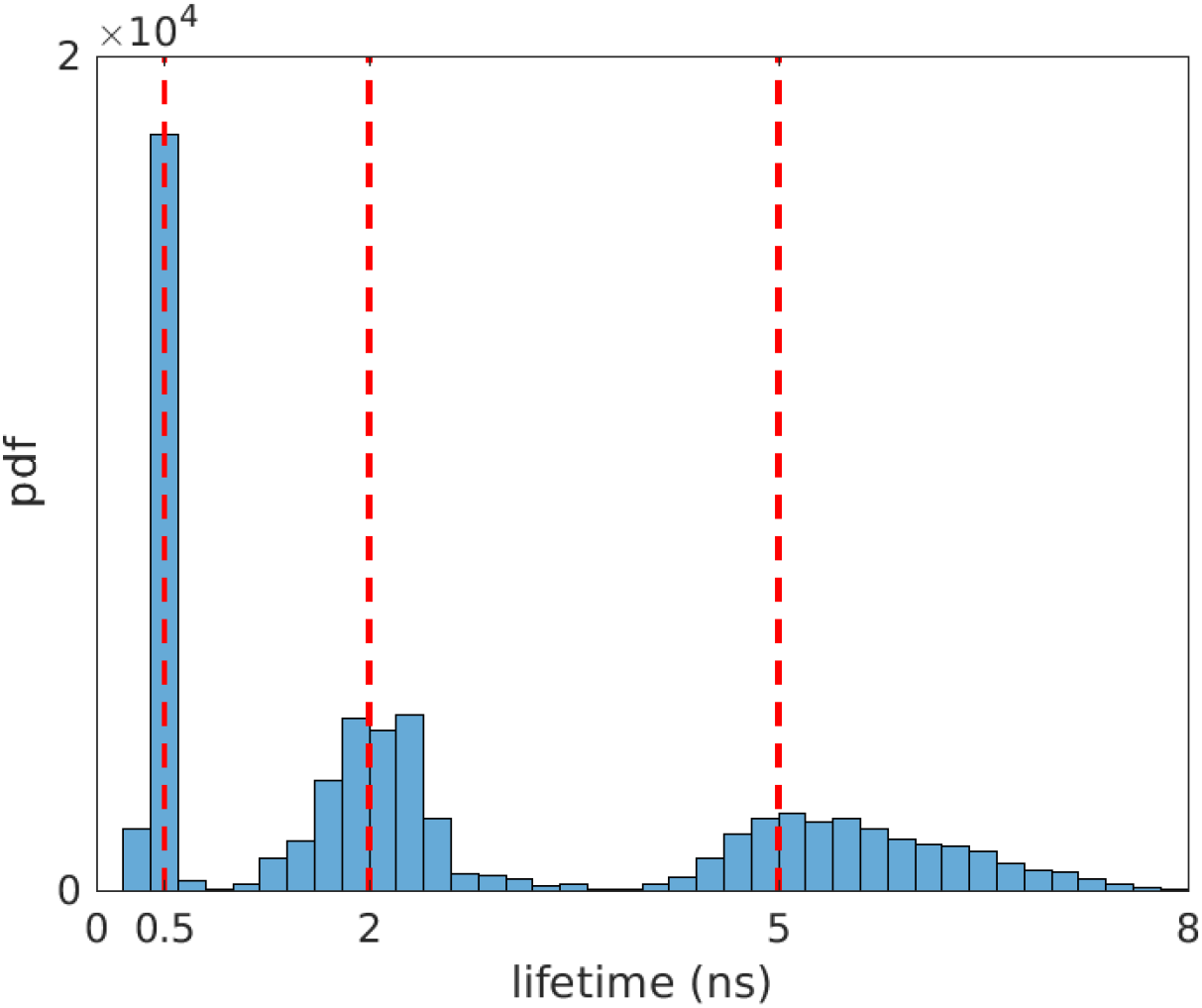
Posterior over 3 lifetimes using 30K synthetic photon arrival times.

Next, we employ experimental data to evaluate the robustness of our method in estimating lifetimes over a wide range, *e*.*g*., short lifetimes falling within the width of the IRF, with short interpulse windows and different photon counts (see Fig. 8). Moreover, employing experimental data containing lifetimes of 0.6 ns, 2.3 ns and 4.6 ns, we will show that BNP-LA can distinguish and deduce 3 lifetime species using as few as 30K photons (see Fig. 9).

**Figure 8:**
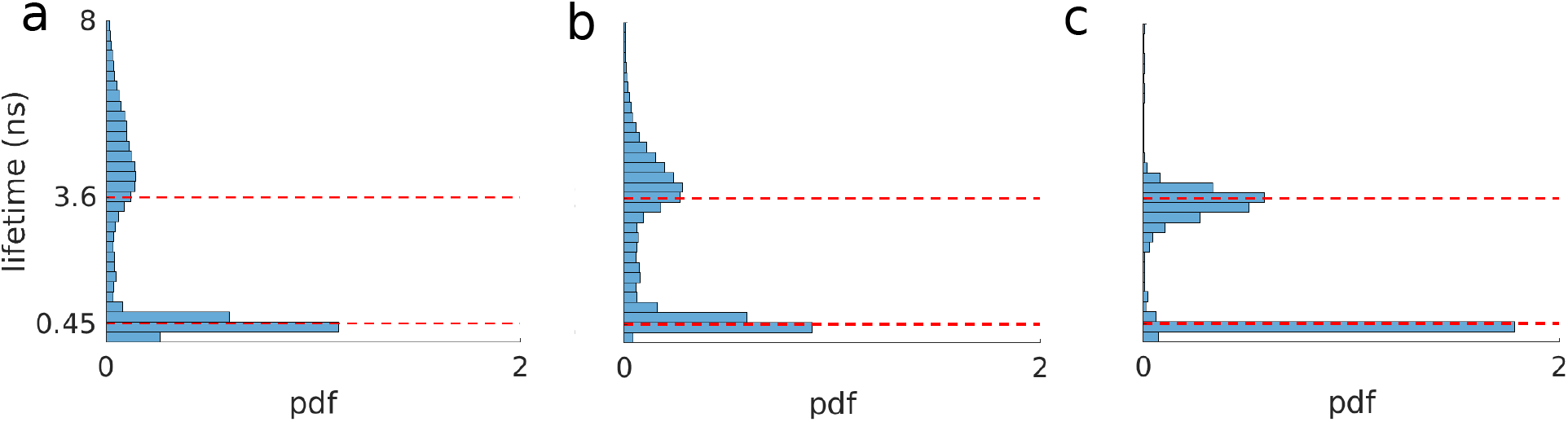
Experimental data with two lifetimes including a lifetime (0.45 ns) below IRF width (0.66 ns). (a-c) Marginal posterior of lifetimes for 500, 1K and 5K photons, respectively.

**Figure 9:**
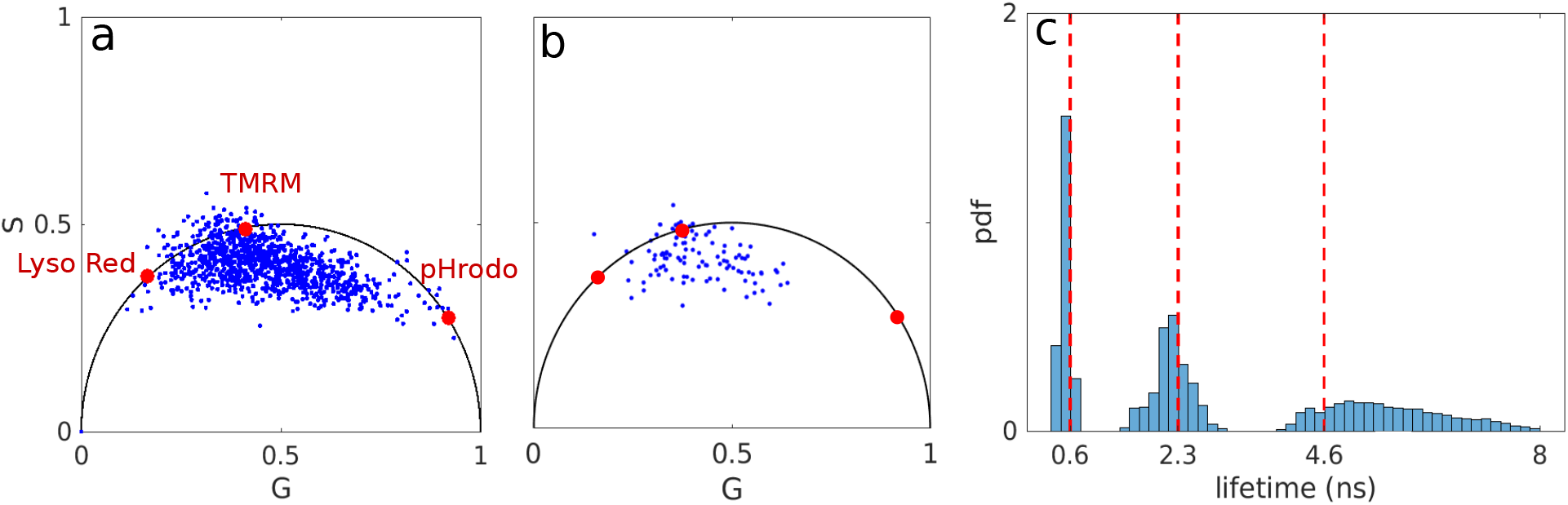
Experimental data containing 3 fluorophore species, namely, Lyso red, TMRM and pHrodo with lifetimes of 4.6 ns, 2.3 ns and 0.6 ns, respectively. (a) For illustrative purposes, we show the results using a phasor plot with 330K photons where the red dots represent the three fluorophore species on the universal circle^26,28^. The corresponding lifetimes are used as ground truth. (b) Results using a phasor plot with 30K photons (which is the same number we analyze). (c) Marginal posterior of lifetimes from the BNP-LA method using 30K photons. The red dashed lines show lifetimes obtained using the phasor technique with 330K photons.

### Synthetic Data

Here, we first illustrate how we simulate our data and then describe our BNP-LA analysis.

To generate synthetic photon arrival time traces, for each photon we would: first need to sample the species being excited; then sample the stochastic excited state lifetime from an exponential; and finally add to this lifetime a stochastic IRF time due to both finite size of laser pulses, *i*.*e*., laser pulses are not infinitely narrow, and detector delay. As such, we first sample the fluorophore species leading to the *k*th photon detection (*s*_*k*_ ∈ {1, …, *M*} for *M* fluorophore species) from a categorical distribution; next excited state lifetime (Δ*t*_em,*k*_) from an exponential distribution; and then add to this the IRF time (Δ*t*_IRF,*k*_) sampled from a normal distribution.

To be clear, the categorical distribution is an extension of the Bernoulli distribution with more than two options (species); the mean of the exponential distribution for each individual species that we use in the simulation to sample lifetimes is set to that species’ lifetime; and the Gaussian used in sampling the IRF time has a mean and standard deviation of 10.4 ns and 0.66 ns (similar to values in our experimental data that we will see shortly)^7^. We also assume a value for the interpulse window of *T* = 12.8 ns again inspired by values from our experimental data.

In cases when the interpulse window is not much larger than both lifetimes and the IRF offset, the data generated as described above can lead to photon arrival times larger than the interpulse window. As such, to guarantee photon arrival times smaller than the interpulse window, we have to introduce a third term as follows

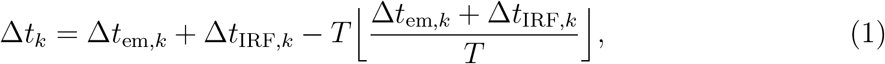

for the *k*th photon arrival time. Here, ⌊⌋ gives the integer part of its argument.

Now, in order to test BNP-LA against different interpulse windows (for which the third term in eq. 1 becomes increasingly important), we simulate data, as described above, using two lifetimes of 1 ns and 8 ns and interpulse windows of 51.2 ns, 25.6 ns, 12.8 ns and 6.4 ns (see Figs. 3a-d).

We start by describing results using the largest interpulse window as this is the easiest case since we can safely ascribe arriving photons as having been generated from the excitation pulse immediately preceding the photon arrival. In this case, the resulting weights ascribed to two lifetimes are non-negligible and add up to more than 0.9. The histogram of the lifetimes pertaining to these wights is shown in Fig. 3e. Here, our method infers both small and large lifetimes with standard deviations of 0.01 and 0.35 ns, respectively. Next, in lower panels f-g, we consider the more difficult case of decreased interpulse windows. This, in turn, leads to larger uncertainties over lifetimes, although the mean of the histograms still coincide with true values. Finally, panel h shows the resulting lifetimes corresponding to the data in panel d with an interpulse window 6.4 ns. Here, again two important weights are found associated to two lifetimes. However, our method begins under-estimating the lifetime (8 ns) larger than the interpulse window. To build an intuition as to why the method begins to fail (as it should) for increasingly small windows, we consider infinitely small interpulse windows. In this case, the photon arrival times are essentially uniform over that window and no information can be extracted from a flat distribution. Conversely, as the interpulse window duration increases, this uniform distribution in arrival times begin acquiring some shape that, loosely speaking, any method can begin leveraging to deduce lifetimes.

Next, we continue by considering the challenging case of multiple lifetimes whose value is smaller than the width of the IRF (in shorthand, “lifetimes below the IRF”). We do so by generating photon traces involving 500, 1K and 5K photons with two lifetimes of 0.2 ns and 0.6 ns, both below the IRF. We then show how many photons are needed to start discerning that we have two lifetimes present. Here, we start by describing the results for the trace containing 500 photons. For this data, while the BNP-LA method ascribes ≈ 0.95 weight to a single lifetime component, the resulting lifetime histogram has a broad range covering both lifetimes (see Fig. 4a). Increasing the photon budget to 1K and 5K, our method begins attributing non-negligible weights to two lifetimes whose sum is larger than 0.9 (see Fig. 4b-c). We also note that the uncertainties over the estimated lifetimes decrease with increasing photon counts.

After demonstrating our method for lifetimes below the IRF, we proceed to assess its performance over a range of lifetimes, namely lifetimes with sub-nanosecond differences (but not necessarily below the IRF), lifetimes comparable to the interpulse window (*T* = 12.8 ns), and lifetimes in the intermediate range. To do so, we analyzed synthetic photon traces containing 500, 1K and 5K photons.

In Figs. 5a-c, we start by considering lifetimes with sub-nanosecond difference (lifetimes of 1 ns and 1.7 ns). Using 500 photons, only a single lifetime is appreciably warranted by the data with a weight much larger than the other lifetimes (see Fig. 5a). However, upon reaching 1K and 5K collected photons, BNP-LA begins ascribing important weight to two lifetimes adding up to more that 0.9; Fig. 5b-c. Next, we examine larger lifetime gaps in Figs. 5d-i. In these cases, the BNP-LA method attributes non-negligible weights to two lifetimes even for datasets with as few as 500 photons; Fig. 5d & g. At larger photon counts, as expected, our method recovers sharper histograms while accurately recovering both lifetimes with less than 8% difference between the histograms’ means (posterios’ means) and ground truth values; Fig. 5e-f & h-i.

Now that we have benchmarked our BNP-LA algorithm using a wide range of lifetimes, we continue by evaluating our algorithm in learning the lifetime weights, designated by *π* ∈ [0, 1]. To be clear, the weights associated to different lifetimes is proportional to the photon ratios from those lifetimes. In order to perform such evaluation, we simulate datasets with two lifetimes of 1 ns and 3 ns containing 1K and 5K photons, the first and second rows in Fig. 6, and weights of (0.5, 0.5), (0.33, 0.66) and (0.2, 0.8) for datasets used in Fig. 6 a-b, c-d and e-f, respectively. As we now see, only weights associated to two lifetimes were found to contribute non-negligibly. Fig. 6 represents histograms over these weights. As expected, cases with 500 photons have larger uncertainty (Fig. 6a-b) with uncertainty decreasing as the photons considered in the analysis mount.

Finally, in Fig. 7, we show that our method is capable of dealing with datasets containing more than two lifetime components. As such, we generate data with three lifetimes of 0.5 ns, 2 ns and 5 ns. These traces are now, naturally, longer (contain more photons) as the inference task is more difficult. In particular, we find that we need about 30K photons to analyze this data though the exact number of photons required is specific to a number of parameters (including how well separated the lifetimes are). The BNP-LA method returned three non-negligible weights adding up to ≈ 0.9 with corresponding mean posterior lifetimes of 0.5 ns, 2 ns and 5 ns. As expected, the resulting lifetime histograms exhibit more uncertainty for larger lifetimes, as measured by a wider posterior, on account of more photon emissions occurring beyond the pulse subsequent to the one resulting in excitation.

### Experimental Data

We now continue by benchmarking our BNP-LA method on experimental data. We start with a dataset acquired using only two fluorophores, namely Calcein and Mito-tracker, with lifetimes of 3.6 ns and 0.45 ns, respectively. Here, the shorter lifetime falls below the IRF, where the IRF parameters as well as the interpulse window are the same as what were used for the simulations. In Figs. 8a-c, we analyze photon traces from this dataset containing 500, 1K and 5K photons, respectively. Our BNP-LA method returns two non-negligible weights adding up to ≈ 0.9 for all the cases. The corresponding lifetime histograms show that the BNP-LA method deduces both lifetimes including the lifetime below the IRF even for as few as 500 photons.

Finally, we test our method employing an experimental dataset containing 3 fluorophore species, *i*.*e*., Lyso red, TMRM and pHrodo, characterized by lifetimes of 4.6 ns, 2.3 ns and 0.6 ns. These “ground truth” values are obtained by using 330K photons with commonly employed phasor plots^26^ (see Fig. 9a).

Next, we process a trace of 30K photons from this data set using both our BNP-LA method and the phasor technique (see Fig. 9b-c). Using only 30K photons, it is difficult to extrapolate the three lifetimes from the phasor plot (Fig. 9b). However, analysis by our method results in three major weights adding up to ≈ 0.9 indicating presence of three lifetime components within the input data. Fig. 9c illustrates the lifetime histogram corresponding to these three weights. The histogram peaks match the values we use as ground truth. Here, for reasons identical to synthetic data, uncertainty increases with larger lifetimes.

## Methods

In this section, we illustrate our likelihood model formulation and inverse strategy. We begin with the likelihood for a set of given photon arrival times 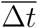

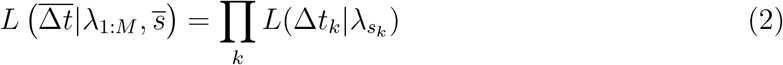

where 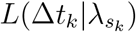 is the likelihood for the *k*th photon arrival time. The indicator parameter *s*_*k*_ allocates photons to different species, 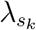 and Δ*t*_*k*_, respectively, denote the inverse of the lifetime and the *k*th photon arrival time. The bars over parameters denote the set of parameters for all *K* photons, for example 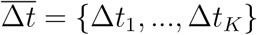.

The likelihood for the *k*th photon arrival time, assuming the photon is from a species indicated by *s*_*k*_, can be derived by considering that Δ*t*_*k*_ is sum of three random variables (also see eq. 1): 1) the time the fluorophore spent in the excited state sampled from an exponential distribution; 2) the stochastic time added due to the IRF sampled from a Gaussian distribution; 3) the number of pulses over which the fluorophore remains excited sampled from a categorical distribution. As such, the likelihood is given by a convolution of the distributions arising from these three contributions^39^

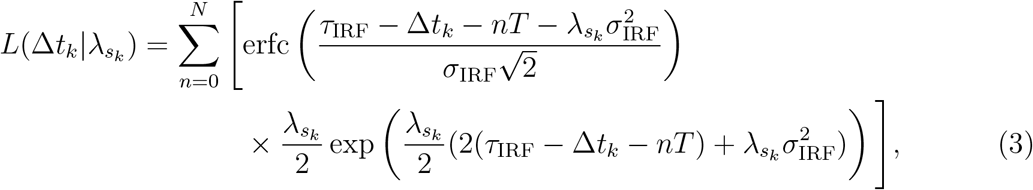

where *N, T, τ*_IRF_ and 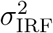 are, respectively, the maximum number of pulses after which photon emissions might occur, the interpulse window, offset, and IRF variance. This likelihood, has been previously derived and employed within a parametric framework with known number of lifetime components^39^. Here, we go beyond the parametric framework and use this likelihood in conjunction with a Dirichlet process to obtain a posterior within a nonparametric Bayesian paradigm where the number of lifetime components is one of the unknowns. We, now, proceed to derive our nonparametric posterior. The posterior is the joint probability over all unknowns we wish to learn including: the weight over each species denoted by symbol *π*_*m*_ for the *m*th species, inverse of the corresponding lifetimes (*i*.*e*., the rate) denoted by *λ*_*m*_ for the *m*th species, and the indicator parameters for each photon designated by *s*_*k*_ and assigning the *k*th photon to one of the species. We collect all these parameters into 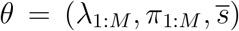 where formally *M* → ∞ within the nonparametric framework. Next, the posterior over *θ*, proportional to the product of the likelihood and priors over these parameters, reads

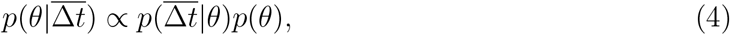

where *p*(*θ*) denotes the corresponding priors. This posterior, however, assumes a nonstandard form and we cannot jointly sample all parameters. Therefore, we invoke a Gibbs sampling strategy for which we can sample individual parameters from the full conditional posteriors^53–60^. That is, the posterior of the parameter of interest conditioned on the remaining parameters. The Gibbs sampling strategy of BNP-LA is as follows:

1. Sample the indicator parameters from their full conditional posterior given by

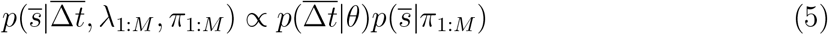

where 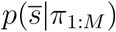 is the prior over the indicator parameters.
2. Sample weights using their corresponding full conditional posterior

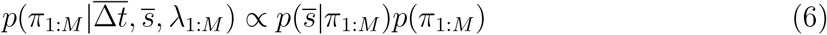

where *p*(*π*_1:*M*_) denotes the prior over weights.
3. Sample the inverse of the lifetimes employing their full conditional posterior given as

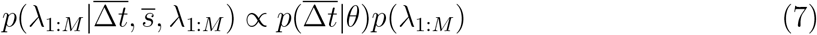

where *p*(*λ*_1:*M*_) is the prior distribution over the lifetime inverse. Now, for the sake of computational convenience, we opt for conjugate priors whenever possible such that those conditional posteriors assume analytical forms allowing for direct sampling. As such, we put a categorical prior distribution over the indicator parameters

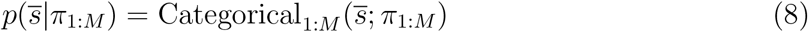

leading to a closed form full conditional distribution that can be directly sampled. For weights, we select a Dirichlet process prior

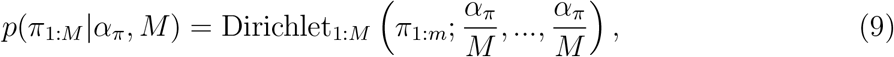

conjugate to the categorical distribution. Here, *α*_*π*_ is a positive hyper-parameter, which we set to one. This results in a standard closed form distribution

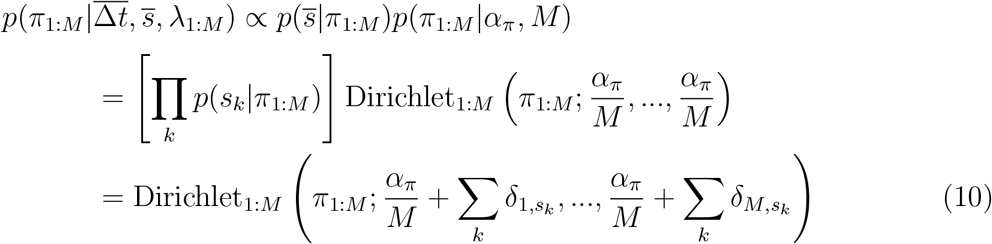

where *δ* denotes the kronecker delta. there are two approaches that can be employed to draw samples from the distribution in eq. 10: slice sampling and finite truncation^46–49^. Here, we opt for finite truncation due its computational efficiency. This approach sets an upper limit on the number of species by assuming a finite but large value for *M* facilitating sampling from the above Dirichlet distribution. Finally, for the inverse of lifetimes, we use a gamma prior to guarantee positive values

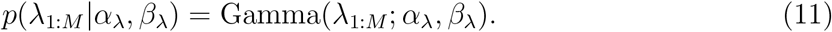

Since the likelihood of eq. 3 does not have an associated conjugate prior, even with a choice of gamma prior, we must use Metropolis-Hastings^61–65^ to numerically draw samples. Here samples are proposed also using a gamma proposal distribution

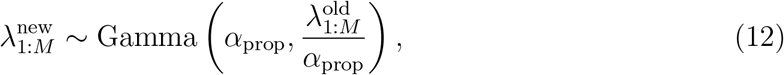

to avoid negative proposals. The proposed values are then accepted with probability

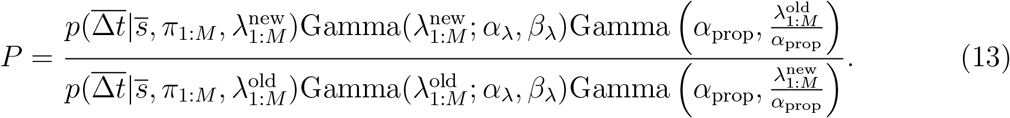

Using the Gibbs sampling strategy described above, we build a chain of samples by iteratively sweeping the set of parameters. Finally, the generated chain can be used for the subsequent numerical analyses.

## Discussion

Fluorescence lifetime experiments provide a means to probe sub-cellular processes and structures. For instance, these techniques have been essential in cancer diagnosis ^66,67^ and monitoring the effect of drugs on cancer cells^67^. However, quantitative assessment of FLIM data remains a challenge as the number of fluorescent lifetime species and their associated lifetimes may vary within a biological sample due to exposure to variable chemical environments^41,42^. These issues immediately require the development of methods capable of learning the number of unique species, as well as their associated lifetimes and photon ratios. Ideally such methods would be robust in treating lifetimes irrespective of what numerical value they ultimately attain in experiments, whether they be shorter than the width of the IRF, on par with the interpulse time, or similar to each other.

Here, we put forward BNP-LA capable of enumerating lifetime components using as few photons as 500 from a single confocal spot while simultaneously deducing the corresponding lifetimes over a wide range from below the IRF to the interpulse window. BNP-LA does so by leveraging tools, such as Dirichlet processes, from the BNP paradigm.

We benchmark BNP-LA using both synthetic and experimental data over a broader range of conditions than was previously accessible. That is, we benchmarked our method against lifetimes shorter than the IRF width, comparable to the interpulse window, and lifetimes with sub-nanosecond gaps with different photon ratios.

In terms of computational cost, the scaling grows is linear with number of photons. While, the exact absolute cost depends on the number of iterations required for the sampler to converge which is, in turn, related to the number of lifetimes and how close they are. For instance,for typical lifetimes in Fig. 8, the computation took approximately 1 minute on a regular scientific desktop. We expect this value to vary depending on the exact CPU specs. Although, here we only assume Gaussian IRFs, BNP-LA can be extended to consider non-Gaussian IRFs by modifications to the likelihood in eq 3. Furthermore, BNP-LA can be used to construct a pixel-by-pixel spatial map of species distributions over a large field of view by independently analyzing data obtained from individual confocal spots across a specimen.

While we have advanced the capability of analysis, there are questions neither we, nor any existing methods can address. Answering these questions, may inspire alternatives to lifetime experiments.

For instance, we cannot learn lifetimes when arrival times far exceed the interpulse window; as a corollary, we cannot avoid posterior broadening for larger lifetimes, *e*.*g*., lifetimes comparable to the interpulse window; we cannot determine whether two exponential components coincide with the same chemical species or two different chemical species; we cannot distinguish chemical species with too small differences, *e*.*g*., as of the current date of publication less than approximately 0.2 ns, with the number of photons we typically analyze (far higher photon counts would then introduce high computational cost which may require greater resources not yet available). Finally, we can only quantify photon ratios, not concentrations nor absorption cross-sections as the latter two quantities always appear as one quantity (a product of both fundamental quantities) in the likelihood.

## Data and Software Availability

The experimental data used in this work are available upon reasonable request from the corresponding author. The software package along with the simulated data used in Fig. 4 are available at https://github.com/MohamadFazel/BNP-LA.

